# Identification of genetic interactions with *priB* links the PriA/PriB DNA replication restart pathway to double-strand DNA break repair in *Escherichia coli*

**DOI:** 10.1101/2022.06.16.496518

**Authors:** Aidan M. McKenzie, Camille Henry, Kevin S. Myers, Michael M. Place, James L. Keck

## Abstract

Collisions between DNA replication complexes (replisomes) and impediments such as damaged DNA or proteins tightly bound to the chromosome lead to premature dissociation of replisomes at least once per cell cycle in *Escherichia coli*. Left unrepaired, these events produce incompletely replicated chromosomes that cannot be properly partitioned into daughter cells. DNA replication restart, the process that reloads replisomes at prematurely terminated sites, is therefore essential in *E. coli* and other bacteria. Three replication restart pathways have been identified in *E. coli*: PriA/PriB, PriA/PriC, and PriC/Rep. A limited number of genetic interactions between replication restart and other genome maintenance pathways have been defined, but a systematic study placing replication restart reactions in a broader cellular context has not been performed. We have utilized transposon insertion sequencing to identify new genetic interactions between DNA replication restart pathways and other cellular systems. Known genetic interactors with the *priB* replication restart gene (uniquely involved in the PriA/PriB pathway) were confirmed and several novel *priB* interactions were discovered. Far fewer connections were found with the PriA/PriC or PriC/Rep pathways, suggesting a primacy role for the PriA/PriB pathway in *E. coli*. Targeted genetic and imaging-based experiments with *priB* and its genetic partners revealed significant double-strand DNA break (DSB) accumulation in strains with mutations in *dam, rep, rdgC, lexA*, or *polA*. Modulating the activity of the RecA recombinase partially suppressed the detrimental effects of *rdgC* or *lexA* mutations in Δ*priB* cells. Taken together, our results highlight roles for several genes in DSB homeostasis and define a genetic network that facilitates DNA repair/processing upstream of PriA/PriB-mediated DNA replication restart in *E. coli*.

**Author Summary:** All organisms rely on DNA replication to grow, develop, and reproduce. In bacteria, the cellular machinery that carries out DNA replication is estimated to fail and prematurely dissociate from the genome at least once per cell cycle. As a result, bacteria have evolved “DNA replication restart” mechanisms that resuscitate failed replication reactions. To probe the function and context of DNA replication restart in the bacterium *Escherichia coli*, we employed a genetic screen to identify genes that were conditionally important in mutant *E. coli* strains compromised in their ability to perform DNA replication restart. Identification of genes with previously known relationships with DNA replication restart confirmed the robustness of our screen, while additional findings implicated novel genetic relationships. Targeted experiments validated the importance of these genes and provided an explanation for their significance in preventing double-strand DNA breaks in cells, a severe form of DNA damage. Our results help to define specific roles for the genes identified by our screen and elucidate the contextual environment of DNA repair upstream of DNA replication restart in *E. coli*.

## Introduction

Cell propagation relies on high-fidelity genome duplication. To accomplish this task, DNA replication complexes (replisomes) loaded onto origins of replication traverse the genome, utilizing parental DNA as templates as they synthesize new DNA strands. During this process, replisomes frequently collide with obstacles such as DNA damage or nucleo-protein complexes [1]. In the most severe instances, these encounters cause replisomes to dissociate from the genome. In *Escherichia coli*, it is estimated that at least once per cell cycle a replisome prematurely dissociates from the chromosome [1, 2]. Bacteria have therefore evolved mechanisms to reload replisomes at premature replication termination sites so that cells can complete genome duplication processes [3, 4].

Genetic and biochemical studies have defined three pathways of DNA replication restart in *E. coli*: PriA/PriB, PriA/PriC, and PriC/Rep (Fig 1) [5, 6]. Null mutations in *priA* or *dnaT* cause similar severe phenotypes, therefore both genes have been placed in the PriA/PriB and PriA/PriC pathways [7-10]. Conversely, minor phenotypes associated with mutations in *priC* or *rep* have placed them in the less frequently utilized PriC/Rep pathway, independent of PriA. The synthetic lethal relationship between *priB* and *priC* aligns with the current three-pathway model as one viable PriA-dependent pathway is maintained with deactivation of either gene (but not both). Additionally, a mutation encoding an ATPase- and helicase-deficient variant of PriA (*priA300*) elicits severe defects when paired with a *priB* deletion, but not a *priC* deletion [11]. Therefore, PriA’s helicase activity is likely required to facilitate the PriA/PriC pathway, but not the PriA/PriB pathway (Fig 1). Each restart pathway recognizes abandoned DNA replication forks, remodels the forks to allow replisome loading, and reloads the replicative helicase (DnaB) with the help of its helicase loader (DnaC). After DnaB is reloaded, it recruits the remaining members of the replisome via protein-protein interactions [12-15].

**Fig 1.**
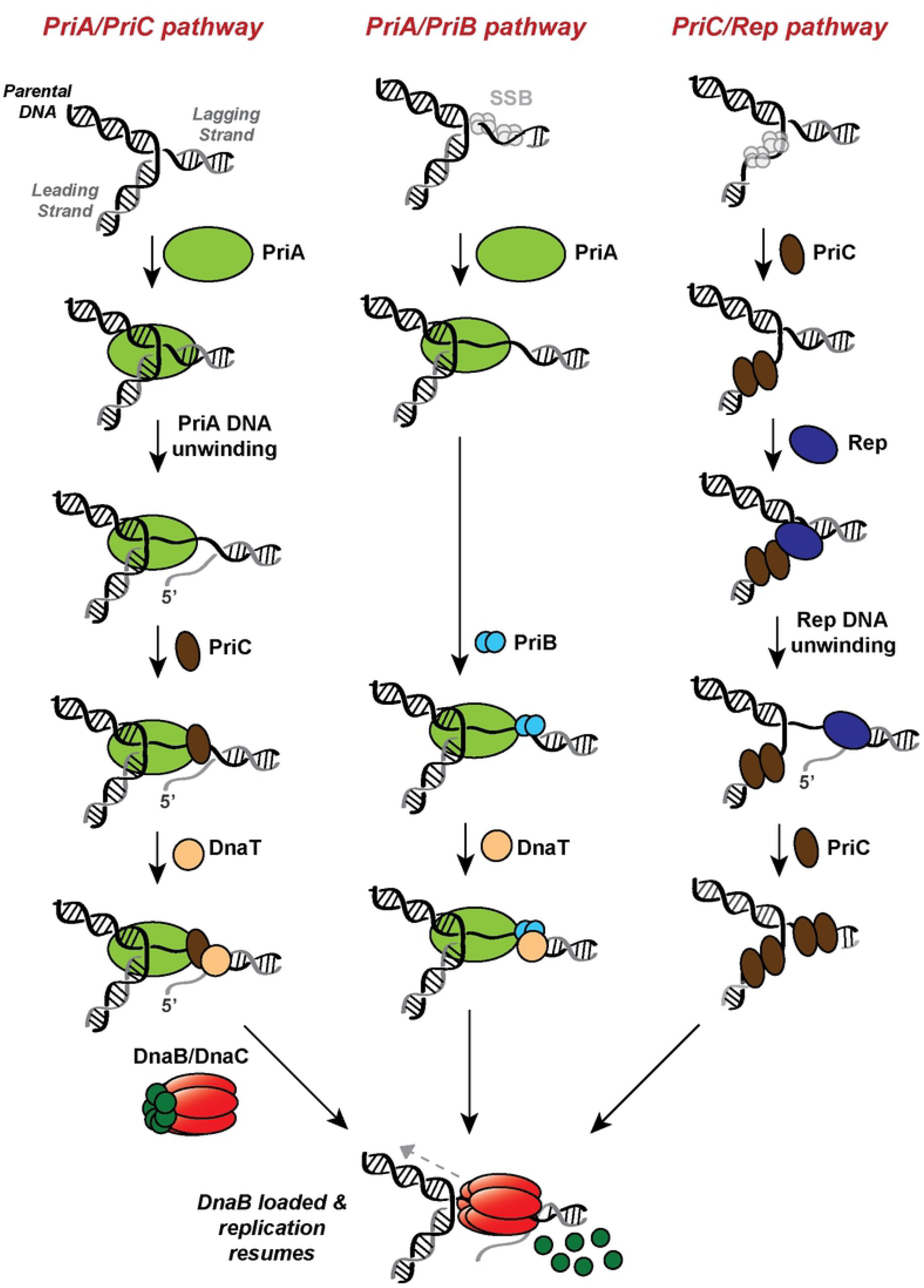
Pathways of DNA replication restart in *E. coli*. PriA/PriC (left) and PriA/PriB (center) pathways efficiently recognize abandoned fork substrates with nascent leading strands, while the PriC/Rep (right) pathway prefers fork substrates with a leading strand gap. All three pathways recognize an abandoned fork, remodel the substrate (if needed) and recruit other replication restart proteins, and load the replicative helicase (DnaB) with the help of the helicase loader (DnaC) to restart DNA replication. The PriA/PriB pathway (center) is inactivated in Δ*priB* cells, the PriA/PriC (left) and PriC/Rep (right) pathways are inactivated in *priC::kan* cells, and the PriA/PriC (left) pathway is inactivated in *priA300* mutants.

Evidence suggests that certain replication restart pathways can be preferentially utilized and/or that each operates on distinct substrates. For example, the PriA/PriB restart pathway appears to be favored following DNA recombination [16]. Mutations in *priB* are also more detrimental than *priC* when paired with a *holD* mutation, which increases instances of fork stalling and collapse [17]. These results could indicate a heavier reliance on PriA/PriB than other pathways for replication restart. Additionally, a *priB* deletion is synthetically lethal with mutations in *dam*, which encodes a DNA methyl transferase whose absence is linked to increased double-strand DNA breaks (DSBs) [18-20]. This observation suggests that PriA/PriB replication restart could be important following DSB repair. Although *priC* disruption alone results in negligible phenotypic effects, *in vitro* evidence suggests that abandoned replication forks with long single-stranded (ss) DNA gaps between the nascent leading strand and parental duplex DNA may be recognized and remodeled efficiently by the PriC/Rep pathway, which could indicate its preference for specific abandoned DNA replication fork structures (Fig 1) [21].

Candidate-based genetic studies have uncovered a limited number of genes linked to DNA replication restart, but a systematic study examining the potential importance of all genes as they relate to this process is lacking. We employed transposon insertion sequencing (*Tn*-seq) in Δ*priB, priC::kan*, and *priA300 E. coli* strains to uncover novel genetic relationships with the DNA replication restart pathways [22-25]. The Δ*priB Tn*-seq screen yielded particularly informative results. The screen and additional genetic experiments corroborated prior genetic results in which *priC, rep*, and *dam* are conditionally essential or important in Δ*priB* cells. Strikingly, the screen also identified many new interactions between *priB* and genes involved in genome maintenance (*lexA, rdgC, uup, rdgB*, and *polA*) and other processes (*nagC*). Mutations in many of these genes produced strong growth defects in Δ*priB* cells, evidenced by plasmid retention, growth competition, and spot plating assays. Furthermore, we found that *rep, lexA, polA*, and *dam* mutants were hypersensitive to ciprofloxacin, which induces DSBs. These mutant strains also accumulated DSBs *in vivo* and displayed significant cell filamentation, a common indicator of poor genomic maintenance. Lastly, some of the toxicity to Δ*priB* cells caused by mutations in *lexA* or *rdgC* appears to result from inappropriate and/or excessive RecA recombinase activity. These results highlight the importance of several genes in Δ*priB E. coli*, define a primary role for the PriA/PriB restart pathway following DSB repair, and help elucidate the interplay between DNA repair and DNA replication restart processes.

## Results

### *Tn*-seq identifies genetic interactions in Δ*priB, priC::kan*, and *priA300* strains

DNA replication restart functionally integrates with other processes in *E. coli*. However, experiments to probe this integration have been limited to candidate genetic and biochemical studies. To systematically map connections between DNA replication restart and other processes, we performed *Tn*-seq screens that assessed the tolerance of gene disruption in mutant strains restricted to specific pathways of DNA replication restart. Current models predict that deleting *priB* inactivates the PriA/PriB pathway, the *priA300* allele (which produces an ATPase- and helicase-deficient PriA variant) disables the PriA/PriC pathway, and a *priC*-null mutation (*priC::kan)* inactivates the PriA/PriC and PriC/Rep pathways (Fig 1) [3, 5, 6, 11]. We therefore carried out screens in each of these backgrounds to independently identify genetic connections with each pathway.

Isogenic wild-type (*wt*), Δ*priB, priA300*, or *priC::kan E. coli* strains were constructed with the *sulB103* mutation, which encodes an FtsZ variant resistant to SulA-mediated cell division inhibition and bolsters the viability of DNA replication restart mutants [26, 27]. Three biological replicate Tn5 transposon libraries with ∼165,000 transposon-insertion mutants were generated for each strain to yield ∼500,000 total insertion mutants in each genetic background. Viable transposon-insertion mutants were selected by plating on Super Optimal Broth (SOB) solid medium supplemented with trimethoprim (ensuring Tn5 insertion). After pooling to assemble each individual replicate, the libraries were subjected to overnight growth in Luria Broth (LB) liquid medium forcing direct competition among transposon-insertion mutants. Successive replication initiation events launch prior to cell division in cells grown in rich media, resulting in more than two replication forks on each chromosome [28-30]. As a result, the sensitivity of the *Tn*-seq screen was likely increased since transposon-insertion mutants were required to facilitate DNA repair and replication restart processes efficiently to prevent collisions between replisomes and unrepaired DNA damage. Following growth in LB, genomic DNA was isolated from each replicate and prepared for next-generation sequencing. The resulting sequencing data revealed the location of transposon insertions as well as relative transposon-insertion mutant abundance. Each gene in our analysis was assigned a normalized weighted read ratio based on insertion tolerance in the mutant strain compared to the *wt* strain [31]. Positive or negative weighted read ratios reflect gene disruptions that were tolerated better or worse, respectively, in the *wt* strain compared to the mutant strain. Genes with few or no insertions were considered important for growth, and such profiles within the *wt* control strain implicated genes as being essential under the tested growth conditions. By comparing insertion profiles of the *wt* and mutant strains, several genes that were conditionally important in replication restart mutant strains were identified.

*Tn-*seq data identified conditional importance for several genes in *E. coli* cells lacking the PriA/PriB restart pathway (Δ*priB*). Genes with the strongest *priB* genetic interactions evidenced by weighted read ratios (Fig 2) and unique insertions (S1A Fig) were selected for subsequent study, except for *rplI* because of its inclusion in the *priB* operon. Corroborating previous studies, the screen implicated *rep* (log_10_ weighted read ratio = 4.11) and *dam* (2.35) as genetic interactors with *priB*. Unexpectedly, *priC* (1.25) was not a prominent hit. Nonetheless, the *priC* gene tolerated no transposon insertions in the Δ*priB* strain and, as described below, the expected lethality of a Δ*priB* Δ*priC* double deletion strain was later confirmed. In addition to known genetic interactions, bioinformatics analysis and manual curation of the *Tn*-seq data implicated a variety of novel gene as genetic interactors with *priB*: *rdgC* (4.06), *nagC* (3.29), *uup* (3.98), *rdgB* (1.86), *polA* (2.80), and *lexA* (2.58) (Fig 2). These top hits (apart from *nagC*) have noted roles in genome maintenance but have not been genetically linked to *priB* prior to this study [32-39]. The abundance of conditionally important genes in Δ*priB* cells is consistent with PriA/PriB serving as the preferred DNA replication restart pathway [17].

**Fig 2.**
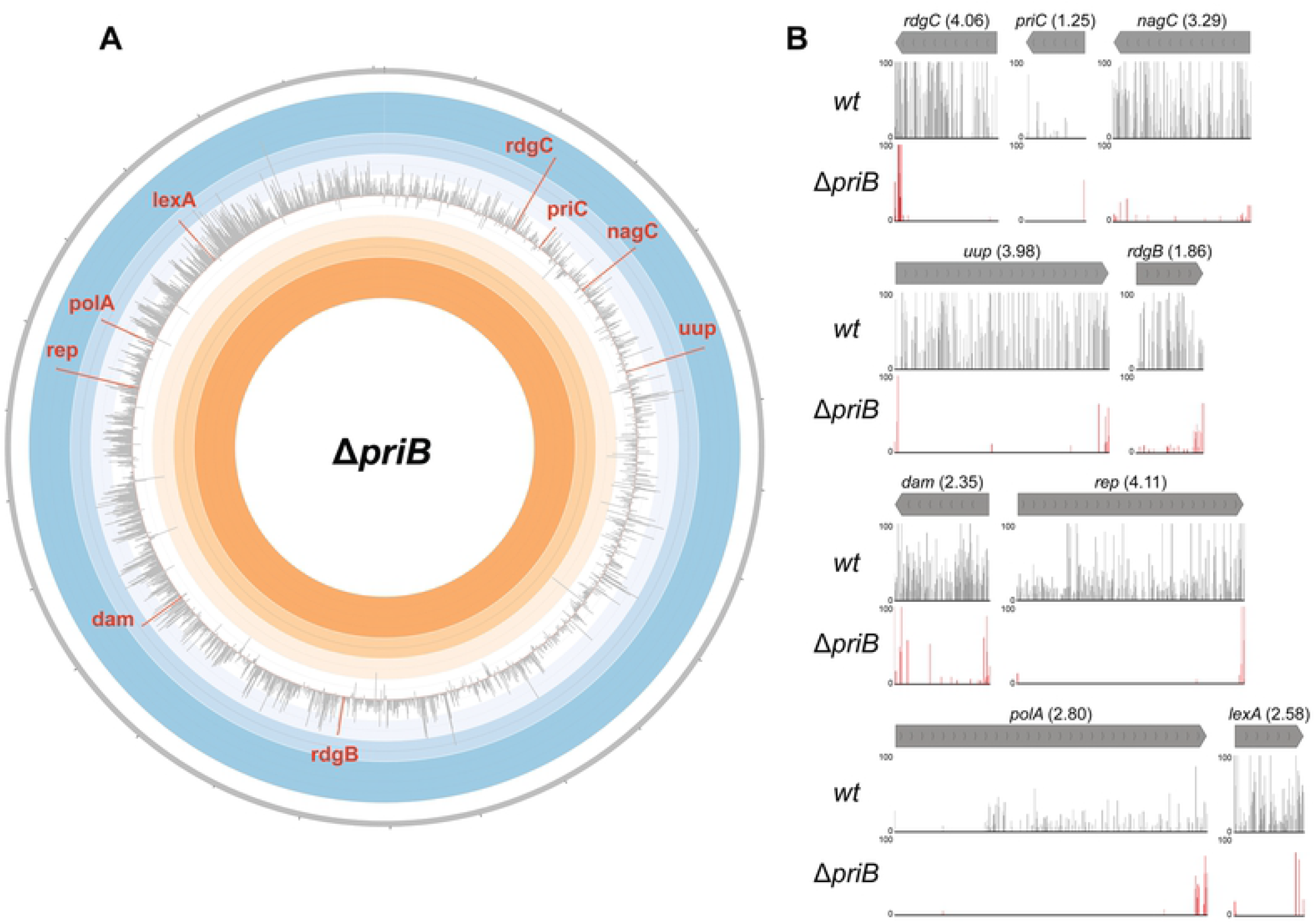
*Tn*-seq results in Δ*priB E. coli*. (A) Circos plot depicting the results of the *Tn*-seq screen in Δ*priB* cells. The effect of single gene disruption via transposon-insertion was determined by comparing *Tn*-seq read profiles in *wt* vs Δ*priB* conditions, yielding a weighted read ratio. Each bar in the Circos plot represents the weighted read ratio (log_10_) of a single gene where extension into the blue or orange region corresponds to a detrimental or beneficial, respectively, effect of gene disruption. Genes with fewer than three unique transposon-insertions per replicate in the *wt* condition are omitted. The individual disruption of many genes involved in genome maintenance produced some of the most prominent defects and were the focus of the study. Bars for notable genes (*rdgC, priC, nagC, uup, rdgB, dam, rep, polA*, and *lexA*) are highlighted in red. (B) MochiView plots for genes highlighted in (A) comparing transposon insertion locations and read abundance. The corresponding weighted read ratio for each gene is included in parentheses. The maximum read height displayed is 100.

Disparities in the transposon-insertion profiles between the *wt* control and *priC::kan* or *priA300* mutant strains were relatively modest, resulting in smaller overall weighted read ratios for genes (S1B,C Fig). This likely was caused by additional mutational stress being tolerated in both mutant strains since each retained the PriA/PriB pathway [3, 5, 6, 11]. One exception was the clear underrepresentation of transposon insertions in *rep* (4.11) within the *priA300* strain (S1C Fig). This result is consistent with the previously described conditional importance of *rep* in *priA300* cells [4, 5, 40]. No other genes were identified with significantly different insertion profiles with respect to weighted read ratios in either the *priC::kan* or *priA300* strains relative to the *wt* control (S1B,C Fig). Interestingly, disruption of the *priB* gene was not significantly less tolerated in the *priC::kan* strain compared to the *wt* strain, but this was due to a very small number of transposon insertions within *priB* for all strains. This is consistent with a prior observation that the *E. coli priB* gene receives fewer insertions in transposition screens than would be predicted for a gene of its size [41].

### Mutations in *priC, rep, lexA, dam, rdgC, uup, nagC*, or *rdgB* confer a dependence on *priB*

Given the importance of the PriA/PriB pathway as reflected by the Δ*priB Tn*-seq screen results, the remainder of our study interrogated the relationship between *priB* and its genetic interactors. A plasmid retention assay was first used to measure the impact of mutations in genes identified in our *Tn*-seq screen on cell viability with or without chromosomal *priB* [42, 43]. The assay followed retention of an unstable, low-copy plasmid (*priB*-pRC7, which contained *priB* and the *lac* operon) in *priB*^+^ or Δ*priB* strains with chromosomal deletions of the *lac* operon and genes identified as conditionally important in the Δ*priB Tn*-seq screen. Plasmid retention or loss was marked by colony color (blue or white, respectively) when plated on SOB-agar containing X-gal and IPTG (Fig 3).

**Fig 3.**
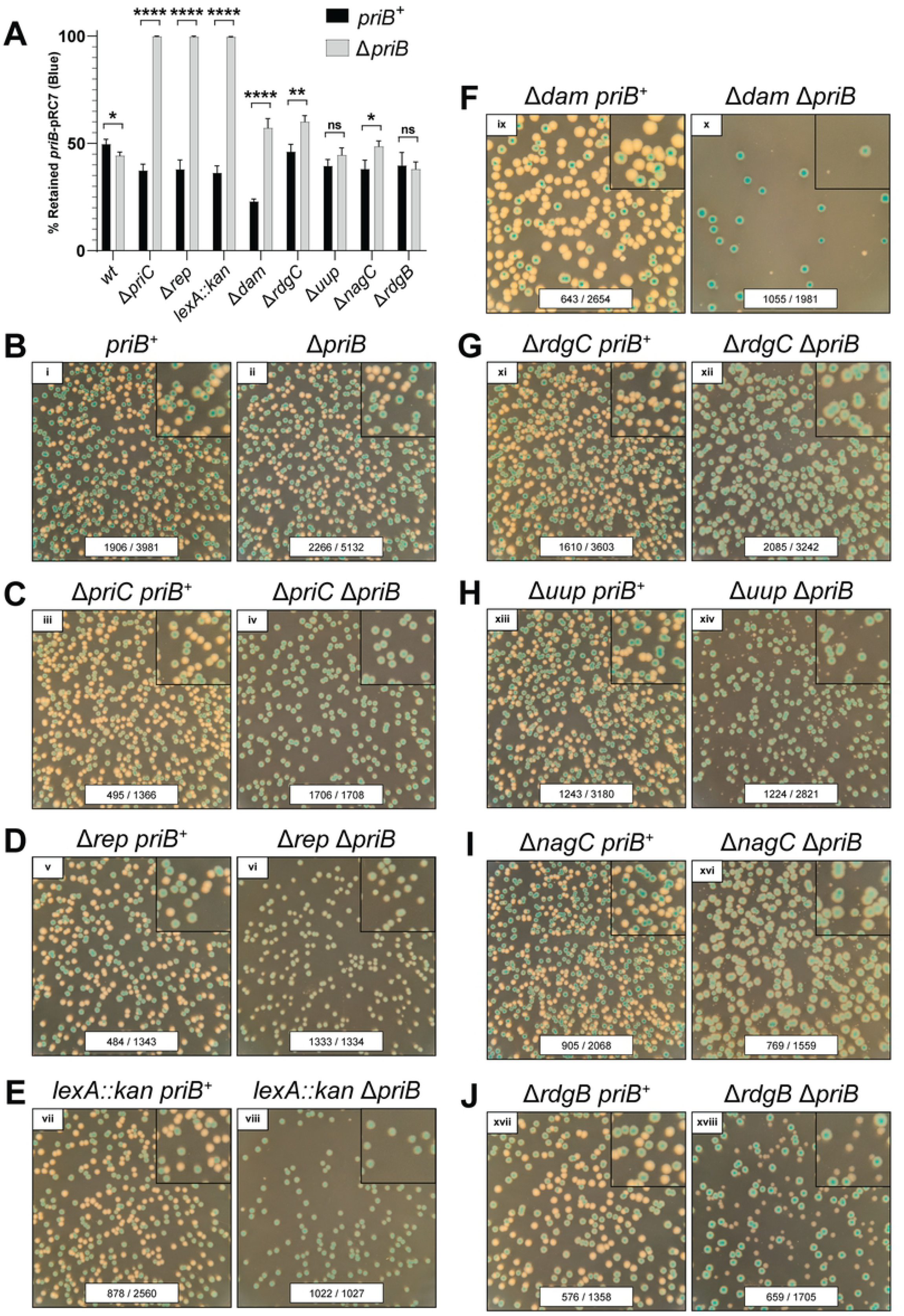
Importance of specific genes in Δ*priB E. coli*. Genes implicated as conditionally important or essential in the Δ*priB Tn*-seq screen were tested with a plasmid retention assay. (A) Percentages of colonies that retained *priB*-pRC7 plasmid are shown. Mean values are depicted with error bars representing standard error of the mean. Statistical significance (unpaired student t-test) for each strain pair is displayed: P < 0.05 (*), P < 0.01 (**), P < 0.001 (***), and P < 0.0001 (****). Representative images from *priB*-pRC7 assay plates are shown as follows: (B) *wt* (C) Δ*priC*, (D) Δ*rep*, (E) *lexA::kan*, (F) Δ*dam*, (G) Δ*rdgC*, (H) Δ*uup*, (I) Δ*nagC*, and (J) Δ*rdgB*. Each plate image includes raw colony counts for each condition (# of blue colonies / # of total colonies). To better visualize small white colonies, 2.25x magnified insets are included in the upper right-hand corner for each plate image. Each plate was incubated at 37 °C for 16 hr.

In line with previous genetic results, deletion of *priC* or *rep* in Δ*priB* cells resulted in persistent retention of *priB*-pRC7, strongly supporting their known synthetic lethal relationships with *priB* (Fig 3C,D) [5]. Screening of a newly identified genetic interaction revealed that *lexA* and *priB* also form a synthetic lethal pair in our genetic background (Fig 3E). LexA is a transcriptional repressor that undergoes auto-proteolysis to induce the SOS DNA-damage response genes [39, 44]. As a result, disruption of *lexA* causes constitutive SOS expression, and it follows that induction of one or more SOS genes is toxic to Δ*priB* cells [27]. For mutations in *priC, rep*, or *lexA*, the extent of plasmid loss was equivalent to control levels in *priB*^+^ cells (Fig 3A-E).

In contrast to the robust and consistent *priB*-pRC7 retention characteristics of the mutant strains described above, mutations in *dam, rdgC, uup, nagC*, or *rdgB* did not prevent plasmid loss when paired with a *priB* deletion (Fig 3F-J). However, white colonies (lacking *priB*-pRC7) formed by these double mutants exhibit reduced growth rates, evidenced by their small size compared to plasmid-containing blue colonies. For Δ*rdgB* Δ*priB*, the disparity in size between blue and white colonies was modest (Fig 3J). However, when other gene deletions (Δ*dam*, Δ*rdgC*, Δ*uup*, or Δ*nagC*) were paired with Δ*priB*, the resulting plasmid-less white colonies were particularly small and difficult to quantify (Fig 3F-I). As a result, disparities in colony size for many of these strains is likely a better proxy of cellular health than a plasmid retention percentage (Fig 3A).

Previous studies have noted a synthetic lethal relationship between *dam* and *priB*, suggesting that DSBs accumulating in Δ*dam* cells are preferentially funneled into the PriA/PriB pathway for restart following their repair [18]. While our data do not confirm a synthetic lethal relationship between *dam* and *priB*, our *priB*-pRC7 retention results strongly support the conditional importance of *dam* in Δ*priB* cells based on a disparity in colony size (Fig 3F). The decreased growth rate of white colonies evident in the assay also confirmed the conditional importance of *rdgC, uup, nagC*, and *rdgB* in Δ*priB* cells (Fig 3G-J). While RdgC (an inhibitor of RecA recombinase activity), Uup (a branched DNA intermediate binding protein), and RdgB (a noncanonical purine pyrophosphatase) have been implicated in genome maintenance processes, these results now map the genes’ interactions with the PriA/PriB restart pathway [32, 35-37, 43, 45]. Surprisingly, conditional importance in Δ*priB* cells extended to *nagC*, which encodes a transcriptional repressor that coordinates *N*-acetylglucosamine biosynthesis but has no known role in DNA metabolism [33, 34]. These data support the notion of conditional importance for *dam, rdgC, uup, nagC*, or *rdgB* in Δ*priB* cells (Fig 3F-J).

### Disruption of *rdgB* in a Δ*priB* strain confers a fitness defect

The disparity of colony sizes in the *priB*-pRC7 retention assay provided only moderate evidence that *rdgB* is conditionally important in Δ*priB* cells. To examine the *rdgB priB* genetic relationship more confidently, the fitness of strains combining *rdgB* and *priB* mutations was tested in a growth competition assay. In this assay, the effect of a *rdgB* deletion was examined within a *priB*^+^ competition (*priB*^+^ vs Δ*rdgB priB*^+^) and within a Δ*priB* competition (Δ*priB* vs Δ*rdgB* Δ*priB*). A synthetic fitness defect would result in selective loss of Δ*rdgB* Δ*priB* in the latter competition. A reporter mutation (Δ*araBAD*) in one strain of each competition was utilized to quantify the relative Δ*rdgB* abundance throughout each competition. As expected for a synthetic *rdgB priB* relationship, simultaneous deletion of both genes caused a pronounced fitness defect within 24 hours when grown in competition with *rdgB*^+^ Δ*priB* cells (Fig 4A, red). In contrast, Δ*rdgB priB*^+^ cells exhibited no detectable fitness defect when grown in competition with *wt* cells, as evidenced by steady relative abundance within the *priB*^+^ competition (Fig 4A, black). These results confirm that *rdgB* is not essential in a Δ*priB* strain, but that it is conditionally important. The mild defect in growth rate of Δ*rdgB* Δ*priB* colonies (Fig 3J) but clear fitness defect (Fig 4A) align well with *rdgB* as a relatively weak hit from our Δ*priB Tn*-seq screen (Figs 2 and S1A), which subjected transposon-insertion mutants to periods of independent and competitive growth.

**Fig 4.**
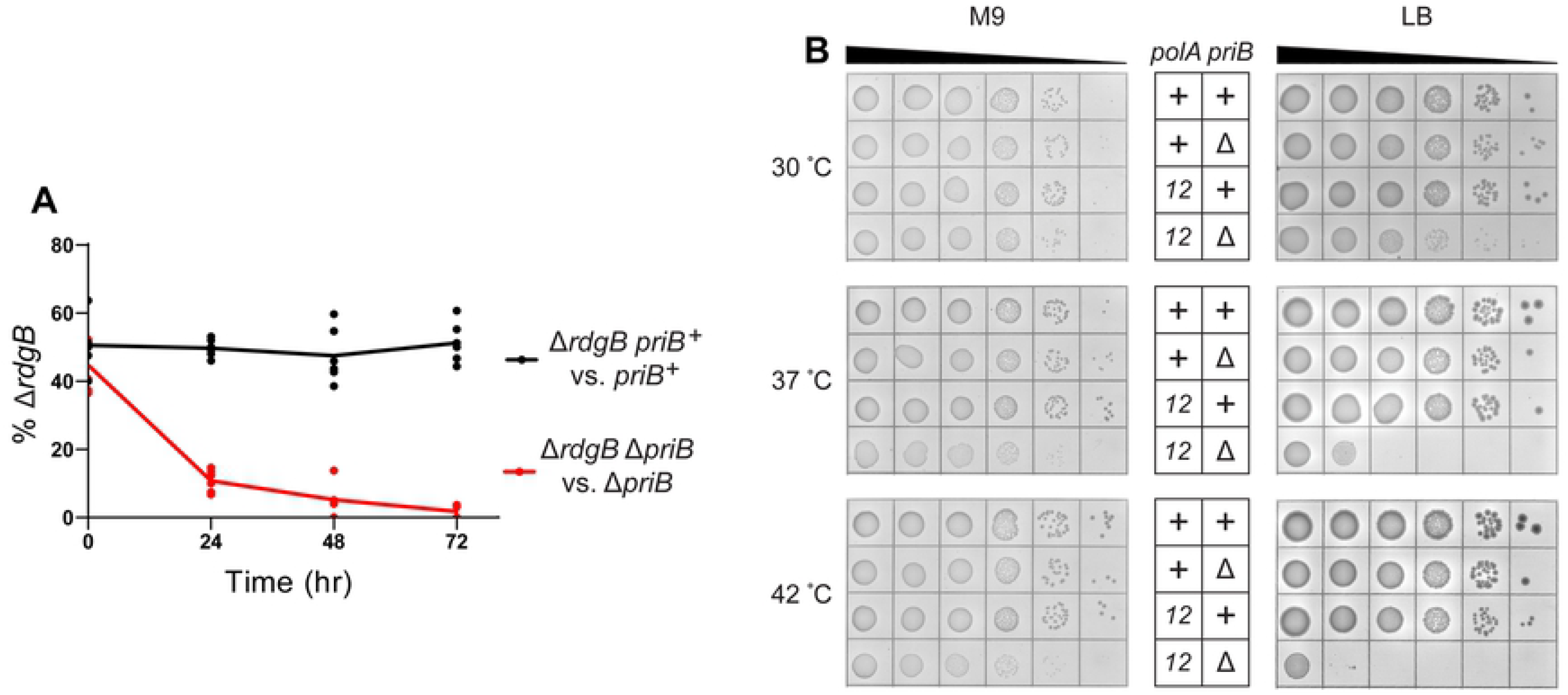
Importance of *rdgB* and Pol I polymerase activity in Δ*priB E. coli*. (A) Growth competition examining the effect of a *rdgB* mutation on fitness *for priB*^+^ or Δ*priB* strains. Trendlines for each series intersect the mean, and biological triplicate data points are presented for competitions done in duplicate. (B) Effect of the *polA12(ts)* allele on *priB*^+^ or Δ*priB* strains. Strains were spot plated on minimal (M9, left) or rich (LB, right) media and incubated at 30, 37, or 42 °C. Dilutions (from left to right) are 10x serial dilutions from normalized overnight cultures.

### Pol I polymerase activity is conditionally important for Δ*priB* cells

The *Tn*-seq screen suggested a genetic relationship between *priB* and *polA* (Fig 2). However, the essential nature of *polA* ruled out simple gene deletion experiments to further examine this link [46, 47]. Inspection of the transposon-insertion profiles (Fig 2B) suggests that only certain regions of *polA* are conditionally important for survival in Δ*priB* cells. Specifically, regions of the gene that encode the C-terminal 3’
s-5’ exonuclease and polymerase domains of DNA polymerase I (Pol I) poorly tolerated transposon insertions in the Δ*priB* strain compared to the *wt* strain. These two domains comprise the Klenow fragment of Pol I [48]. Conversely, the portion of *polA* encoding the 5’-3’ exonuclease domain poorly tolerated insertions in both the Δ*priB* and *wt* strains, consistent with this domain encoding the essential function of *polA* in rich media [46].

To test the importance of the polymerase activity of Pol I in Δ*priB* cells, we utilized the *polA12(ts)* mutant allele. This mutation encodes a Pol I variant with severely inhibited polymerase activity at high temperatures [49-51]. Additionally, *polA12(ts)* is synthetically lethal with a *priA* mutation under non-permissive conditions [7, 52]. Spot plate assays examined the viability of *polA12(ts)* Δ*priB* and control strains at increasing temperatures on LB (rich) or M9 (minimal) media to determine the conditional importance of Pol I polymerase activity (Fig 4B). In agreement with the *Tn*-seq screen results, *polA12(ts)* Δ*priB* cells displayed temperature-sensitive synthetic defects on LB media. At 37 °C, the double mutant was at least 100-fold less viable than the *polA12(ts) priB*^*+*^ strain, and this effect was exacerbated to ∼1000-fold at 42 °C. The *polA12(ts)* mutation appeared to cause a reduced growth rate of Δ*priB* cells even at 30 °C, evidenced by the smaller colony sizes in the double mutant. Based on previous studies, this detrimental effect is likely driven by reduced polymerase activity [49-51]. Interestingly, *polA12(ts)* Δ*priB* strain viability was significantly restored by plating on M9 (minimal) media. This partial suppression likely stems from tighter control over DNA replication initiation in minimal media, and it underpins the importance of efficient genome maintenance in nutrient-rich environments [28-30].

### Mutations in *rep, lexA, polA*, or *dam* cause sensitivity to exogenous DSBs

A prior study demonstrated a synthetic lethal relationship between *priB* and *dam*, and suggested that this relationship may result from DSBs formed in *dam* mutants being funneled into the PriA/PriB restart pathway following their repair [18]. Therefore, we examined whether other genes identified by our Δ*priB Tn*-seq screen could be driving toxicity through enhanced DSB accumulation. Mutant strains were spot plated onto LB medium supplemented with sublethal concentrations of the DSB-inducing antibiotic ciprofloxacin (Fig 5) [53, 54]. A *recA* deletion strain was utilized as a positive control for hypersensitivity [55] and was inviable at 5 ng/mL ciprofloxacin. Notably, a Δ*priB* strain also exhibited extreme hypersensitivity and was inviable at 10 ng/mL ciprofloxacin. Mutations in *rep* and *lexA* led to viability defects at 10 ng/mL ciprofloxacin but were significantly more resistant than Δ*priB* or Δ*recA* strains. At 15 ng/mL ciprofloxacin, the Δ*dam* and *polA12(ts)* mutants began to display defects as well. Other mutants identified in the Δ*priB Tn*-seq screen were not sensitized to ciprofloxacin (S2 Fig). These results suggest cellular roles for *priB, recA, dam, rep, lexA*, and *polA* in prevention and/or repair of DSBs *in vivo*.

**Fig 5.**
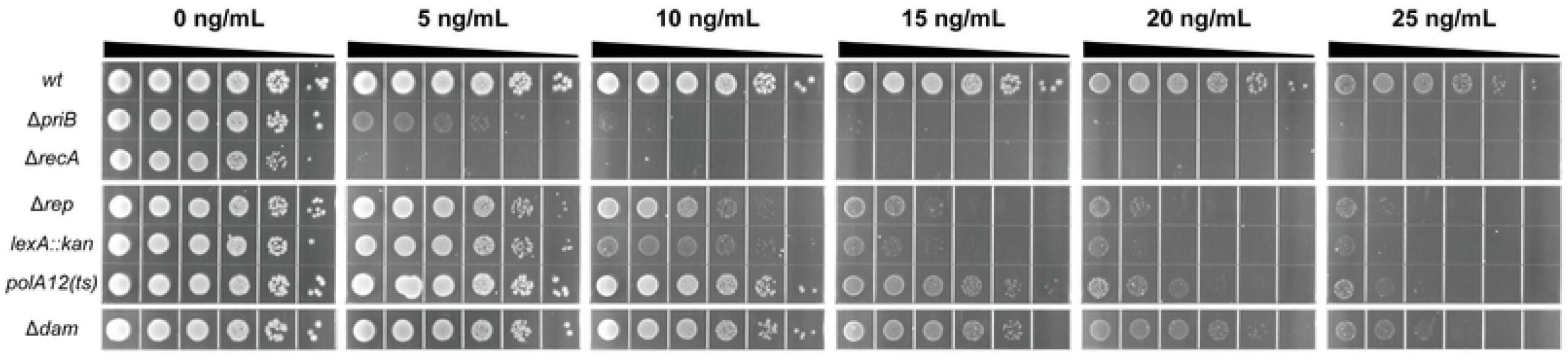
Effects of *priB, rep, lexA, polA*, or *dam* mutations on DNA damage sensitivity in *E. coli*. Sensitivity of mutants to DSBs was examined by spot plating on LB-agar with 0-25 ng/mL ciprofloxacin. A *recA* deletion strain was utilized as a positive control of ciprofloxacin hypersensitivity. Dilutions (from left to right) are 10x serial dilutions from normalized overnight culture. Displayed spot plate data are representative of three replicates.

### Visualizing DSBs *in vivo* with MuGam-GFP

Sensitization of *rep, lexA, polA*, and *dam* mutants to ciprofloxacin suggests that these mutant strains may also have enhanced levels of endogenous DSBs. To test this hypothesis, mutations were transduced into an *E. coli* strain (SMR14334) encoding inducible MuGam-GFP, a DSB sensor protein, and the extent of DSB accumulation was determined *in vivo* with fluorescence microscopy (S3A,B,D Fig) [56].

MuGam-GFP foci were more abundant in a *dam* deletion strain than in the *wt* strain (Fig 6A,B) [20]. These mutant cells were also severely filamented which is a hallmark of genome instability in *E. coli* (Fig 6A,C) [57]. Consistent with their sensitivity to ciprofloxacin (Fig 5), mutations in *rep, lexA*, or *polA* also resulted in increased MuGam-GFP focus formation (Fig 6B) and cell length (Fig 6C). Notably, a *rdgC* mutant displayed significant accumulation of DSBs (S3A,B,D Fig) while exhibiting only a moderate increase in cell length (S3C Fig) and no observable sensitization to ciprofloxacin (S2 Fig). It is worth noting that MuGam helps repair DSBs via non-homologous end-joining, therefore, the extent of DSB formation may be more dramatic in these strains than the results presented here [58].

**Fig 6.**
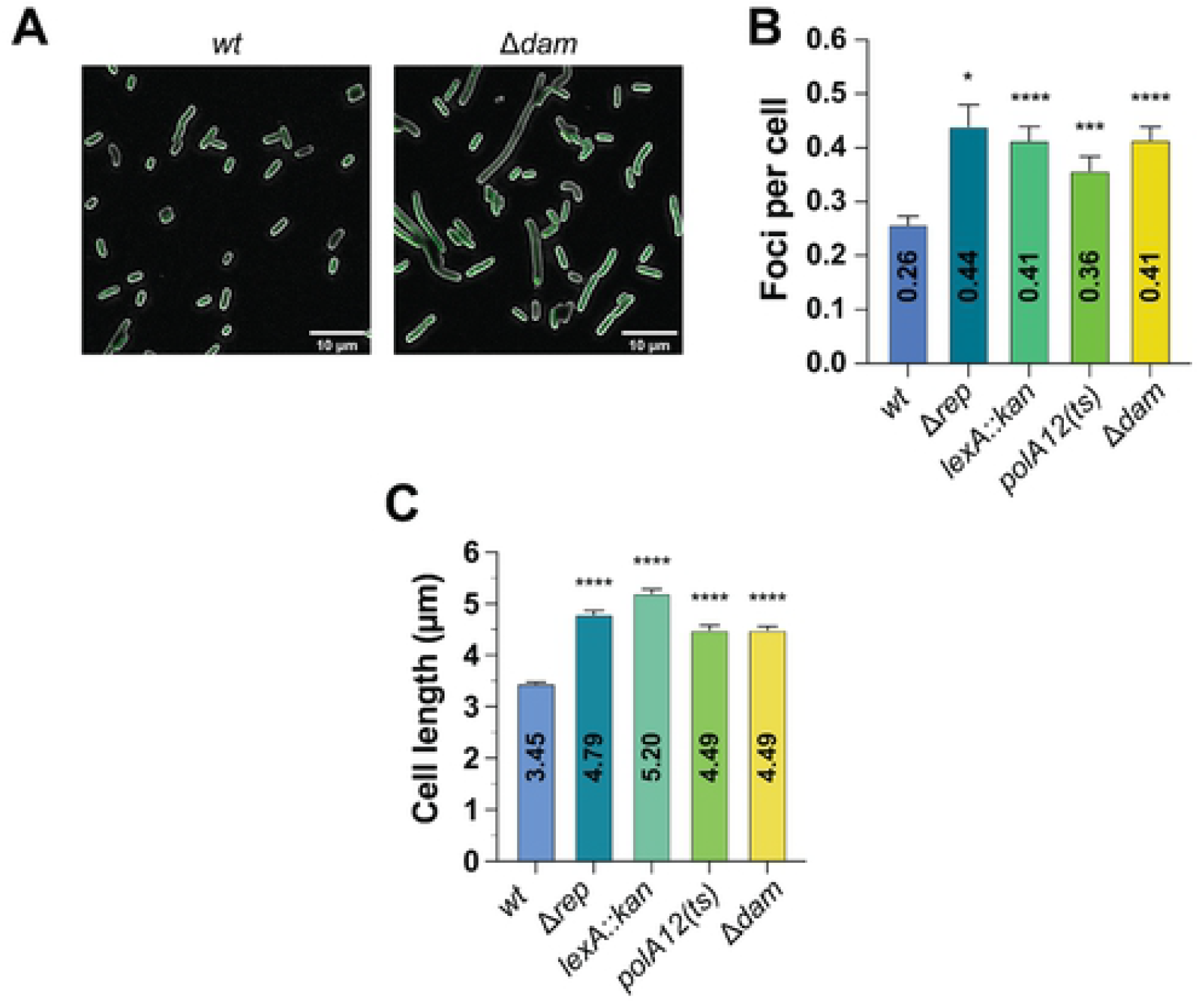
Enhanced DSB formation in mutant *E. coli* strains. (A) Representative images depicting MuGam-GFP (green) foci and FM 4-64-stained membranes (gray) for *wt* (left) and Δ*dam* (right) strains. The abundance of MuGam-GFP foci per cell (B) and measured cell lengths (C) are displayed for *wt*, Δ*rep, lexA::kan, polA12(ts)*, and Δ*dam* strains. (B,C) Mean values are depicted with error bars representing standard error of the mean. Statistical significance (U-Mann-Whitney) for each strain compared to the *wt* control is displayed: P < 0.05 (*), P < 0.01 (**), P < 0.001 (***), and P < 0.0001 (****).

The evidence of DSB accumulation and cell filamentation in other mutants tested is less compelling. Mutations in *priC, uup*, or *rdgB* produce only mild filamentation phenotypes, and there was limited evidence that disrupting *priC* enhances DSB levels (S3A-D Fig). In fact, *nagC* and *rdgB* mutant strains exhibited significantly lower abundance of MuGam-GFP foci compared to the *wt* control and GFP focus levels in the *nagC* mutant approached the lower limit of detection. For the *nagC* mutant, this may have been caused by a significantly lower level of mean fluorescence (S3E Fig).

### Modulating RecA function partially suppresses *lexA* or *rdgC* mutational effects on Δ*priB* cells

Mutations in *dam, rep, lexA, polA*, or *rdgC* increase DSB formation *in vivo* (Figs 6A,B and S3A,B,D). In most cases, this effect is accompanied by sensitization to ciprofloxacin (Figs 5 and S2) and cell filamentation (Figs 6C and S3A,C). Deleting *dam* or hindering Pol I polymerase activity can cause persistent ssDNA gaps that form DSBs when subsequent replisomes collide [59-61]. Similarly, a loss of Rep accessory helicase activity correlates with more stalled replication forks that can create DSBs when they are encountered by subsequent replisomes [62, 63]. Our data strongly suggest an increase in DSB formation in *lexA* or *rdgC* mutants, which likely accounts for their genetic relationships with *priB*, but their mode of DSB formation is less clear.

Previous work has shown that loss of PriA or Rep helicase activity at stalled replication forks can cause inappropriate RecA recombinase loading mediated by the ssDNA gap repair proteins RecFOR [40]. After it is loaded by RecFOR, RecA is hypothesized to reverse a stalled replication fork to form a Holliday junction, also known as a “chicken-foot” structure [64, 65]. Because LexA or RdgC inhibit the activity of cellular RecA (via transcriptional repression [39] or physical inhibition [32], respectively), we hypothesized that more stalled forks were reversed in *lexA* or *rdgC* mutants. The DSBs observed *in vivo* (Figs 6B and S3A,B) could form in these mutants when the “chicken-foot” structures were encountered by additional replisomes (from multi-fork replication conditions in rich media) or upon processing by RuvABC, the Holliday junction resolvase [28, 63, 66].

To test this hypothesis, we examined the effect of RecA modulation on *lexA* or *rdgC* mutants in the *priB*-pRC7 plasmid retention assay (Fig 7). Previously, our results identified a conditional essentiality of *lexA* in Δ*priB* cells based on robust retention of the *priB*-pRC7 plasmid (Figs 3E and 7A,B). After deleting *recR* in this strain (inactivating the RecFOR pathway), we observed viable *lexA::kan* Δ*priB* white colonies (Fig 7C). The resulting colonies display considerable growth defects, but these results strongly support a partial suppression of *lexA::kan* Δ*priB* via *recR* deletion. Likewise, the conditional importance of *rdgC* in Δ*priB* cells (Fig 3G) was partially suppressed with a *recR* deletion, evidenced by significantly larger plasmid-less white colonies (Fig 7C).

**Fig 7.**
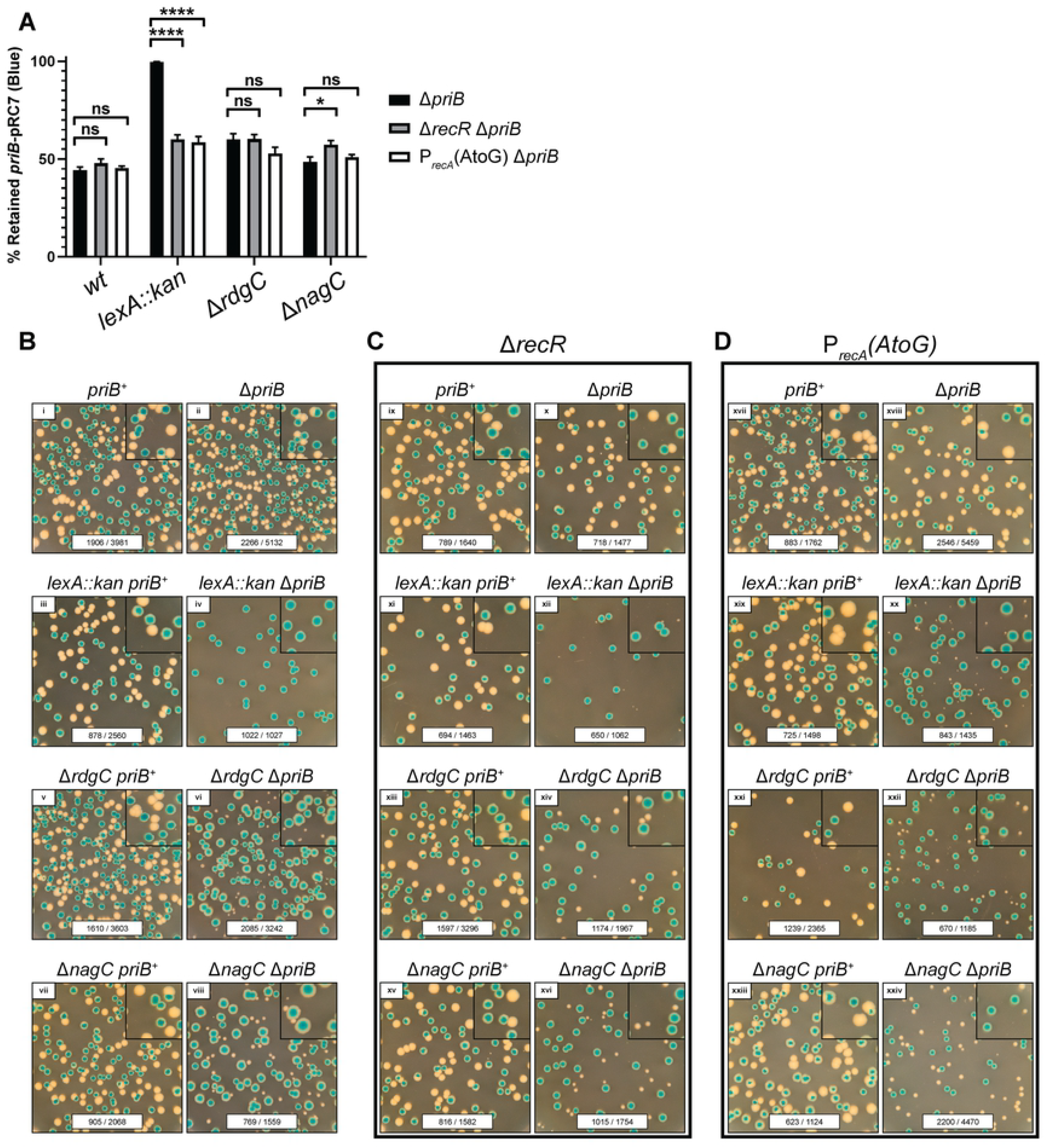
Modulating RecA activity partially suppresses mutational effects on Δ*priB E. coli*. (A) Plasmid (*priB*-pRC7) retention in Δ*priB* strains is shown along with strains also carrying a *recR* deletion or *recA* promoter mutation. Mean values are depicted with error bars representing standard error of the mean. Statistical significance (unpaired student t-test) for each strain pair is displayed: P < 0.05 (*), P < 0.01 (**), P < 0.001 (***), and P < 0.0001 (****). (B) Representative images of *priB*-pRC7 assay plates are presented for *wt, lexA::kan*, Δ*rdgC*, and Δ*nagC* strains with or without chromosomal *priB*. This experiment was extended to strains with (C) a *recR* deletion or (D) a mutation in *recA*’s promoter. (B-D) Each image includes raw colony counts for each condition (# of blue colonies / # of total colonies). To better visualize small white colonies, 2.25x magnified insets are included in the upper right-hand corner. Each plate was incubated at 37 °C for 22 hr.

In addition to restricting the scope of RecA activity *in vivo* with a *recR* deletion, we hypothesized that reducing the cellular levels of RecA would also produce a suppressive effect. To accomplish this, we utilized a *recA* promoter mutation, P_*recA*_(AtoG), which decreases *recA* expression [43, 67, 68]. This mutation also suppressed the effects of *lexA* or *rdgC* mutations in Δ*priB* cells, and the degree of suppression was strikingly similar to that of a *recR* deletion (Fig 7D). To rule out general suppression ability of these RecA modulations, we tested their effect on other mutants identified in our *Tn*-seq screen. We only observed modest evidence of suppression by RecA modulation in Δ*nagC* Δ*priB* strains when comparing the relative sizes of white and blue colonies (Fig 7B-D). Taken together, these results suggest that *lexA* or *rdgC* deletions promote inappropriate and/or excessive RecA activity causing stalled replication forks to physically reverse and eventually devolve to DSBs upon replisome collision or Holliday junction processing.

## Discussion

DNA replication restart reactivates prematurely abandoned DNA replication sites that have failed due to replisome encounters with damaged DNA or proteins tightly bound to chromosomes. Our knowledge of the coordination between DNA replication restart and other genome maintenance pathways has been limited by a lack of systematic genetic studies assessing the importance of genes to each replication restart pathway in *E. coli*. To determine links between replication restart and other cellular processes, we have identified genes that are conditionally essential or important in *E. coli* strains with inactivated replication restart pathways. High-density transposon mutant libraries in strains lacking *priB, priC*, or with the *priA300* mutation were analyzed after growth on rich media. These mutations inactivate the PriA/PriB, PriC/Rep and PriA/PriC, or PriA/PriC pathways, respectively (Fig 1) [5]. Comparison of transposon insertion profiles to a *wt* control strain revealed genetic interactions with specific replication restart pathways.

Several genes were found to be conditionally essential or important in Δ*priB E. coli*, which specifically lacks the PriA/PriB pathway (Figs 2 and S1A). In contrast, only one gene (*rep*) displayed significant importance in *priA300 E. coli* and no genes were significantly conditionally important in *priC::kan E. coli* (S1B,C Fig). These results point to PriA/PriB serving as the major replication restart pathway integrated within the larger genome maintenance program in *E. coli*, consistent with prior data [17]. It is possible that the PriA/PriC and PriC/Rep pathways operate on DNA replication fork substrates that are rarely generated under the conditions tested in our experiments.

Deletion of *rep* was found to be detrimental in both Δ*priB* and *priA300* strains, consistent with a general importance of the Rep helicase in genome maintenance (Figs 2 and S1C). Rep can be recruited to stalled replication forks via interaction with PriC where it helps facilitate DNA replication restart in the PriC/Rep pathway (Fig 1) [69, 70]. PriC interaction with Rep also stimulates its helicase activity [71]. It may be that Δ*priB* and *priA300 E. coli* strains rely more heavily on the PriC/Rep pathway or that deletion of *rep* places a larger burden on the PriA/PriB or PriA/PriC DNA replication pathways. In accordance with the latter possibility, Rep also interacts with the replicative helicase, DnaB, which localizes Rep helicase activity to sites of DNA replication and is thought to enhance its ability to remove tightly associated protein barriers ahead of the replication fork [70]. The absence of Rep results in increased fork stalling, replisome dissociation, and DSBs if left unrepaired, which could also feed into the PriA/PriB pathway (Fig 8) [62, 63].

**Fig 8.**
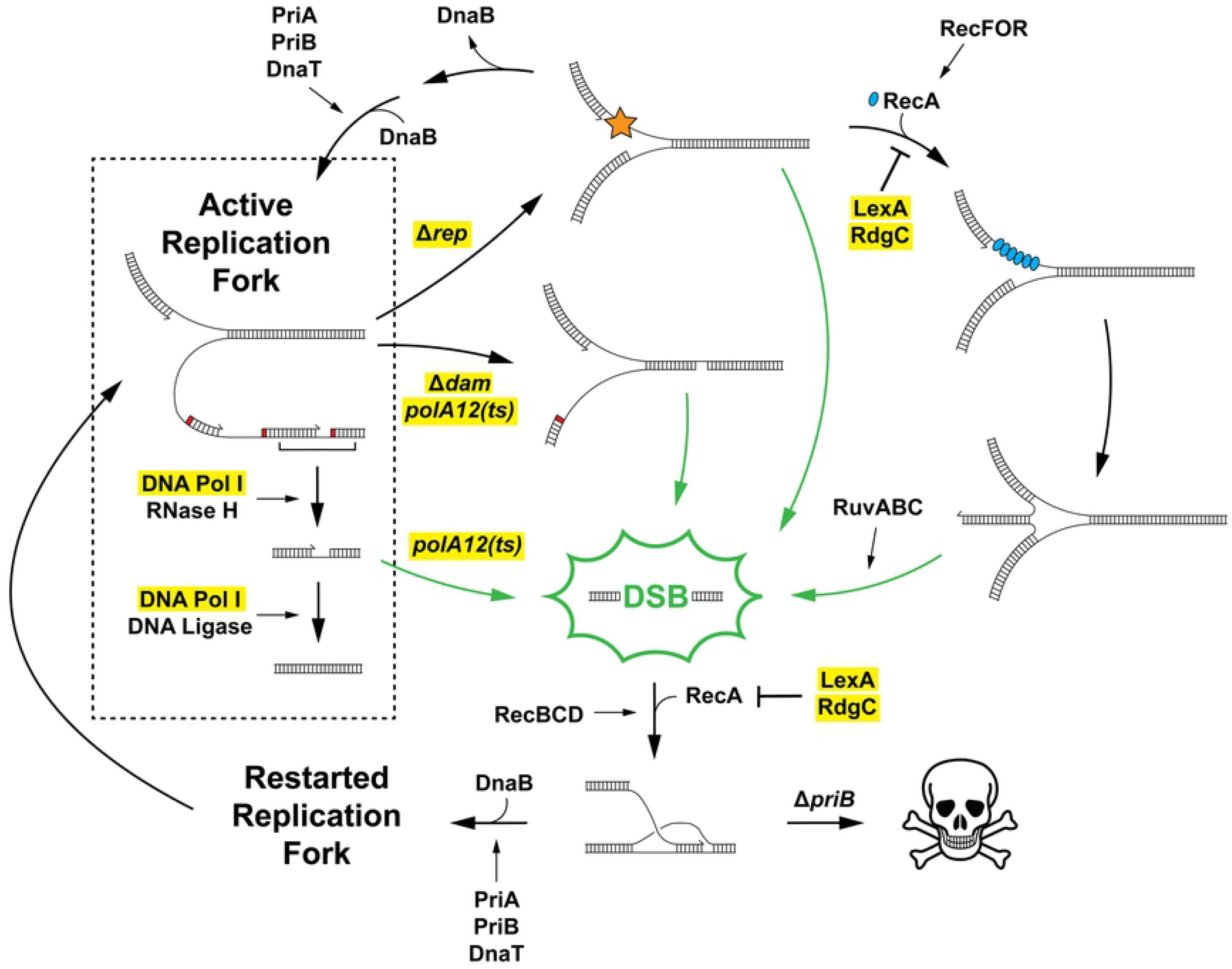
DSBs accumulate from a variety of sources and are funneled into the PriA/PriB replication restart pathway following their repair. An active replication fork facilitates continuous DNA synthesis on the leading strand, while lagging strand synthesis is discontinuous and downstream processing is required by other enzymes. These productive processes are contained within the black box. Several damaging paths are also shown. Loss of Rep causes an increase in replication fork collisions with nucleo-protein complexes (orange star). The most severe collisions cause lethal replisome dissociation unless DNA replication restart is carried out, which is primarily facilitated by the PriA/PriB pathway. Increased mismatch repair (without Dam methylation) or loss of Pol I polymerase activity following DNA repair or during Okazaki fragment maturation cause persistent ssDNA gaps. RecA (blue) loading at stalled replication forks mediated by RecFOR can drive fork reversal, which is inhibited by LexA or RdgC. Stalled/reversed replication forks and ssDNA gaps are DSB-prone substrates; if they are not efficiently repaired, they lead to DSBs (green arrows) when they are encountered by subsequent replisomes. When DSBs form, they are recognized and repaired with homologous recombination (RecA is loaded via RecBCD pathway). The resulting D-loop substrate is shuttled into the PriA/PriB pathway to reinitiate DNA replication and maintain cell viability. The genes/proteins examined in this study are highlighted in yellow.

In addition to the known importance of *rep* in Δ*priB* cells, our results corroborated the importance of *dam* and *priC* in Δ*priB* cells (Fig 3C,F) [5, 18]. In cells lacking Dam methyltransferase, both DNA strands are nicked and excised at equal frequency by methyl-directed mismatch repair enzymes, causing persistent ssDNA gaps that can lead to DSBs (Fig 8) [59]. Interestingly, Δ*dam* cells are also associated with chromosomal over-replication, likely stemming from DSB repair feeding into DNA replication restart [72]. Over-replication could exacerbate DSB accumulation in Δ*dam* cells and it may elicit a similar effect in other DSB-causing mutants described in this study. The synthetic lethality of the Δ*priB* Δ*priC* combination was also confirmed (Fig 3C), although the genetic relationship was not detected in either the Δ*priB* or *priC::kan Tn*-seq screens due to a small number of transpositions insertions mapped for *priB* or *priC* in the *wt* reference strain (Figs 2 and S1B). This may be due to a transposition recalcitrance for *priC* as has been noted for *priB* [41]. Thus, it is possible that additional *priB, priC*, or *priA300* genetic interactions beyond those described here may exist and that limitations of the *Tn*-seq approach could mask their identification.

The *Tn*-seq results in the Δ*priB* strain and targeted genetic experiments identified a host of novel *priB* genetic interactors: *lexA, polA, rdgC, uup, nagC*, and *rdgB* (Figs 2 and 3). In addition to mutant strains expected to exhibit DSB accumulation (*rep* and *dam*), *in vivo* measurements detected significant DSB accumulation for *lexA, polA*, and *rdgC* mutants (Figs 6A,B and S3A,B). Formation of DSBs in these mutant strains was correlated with longer cell lengths (Figs 6C and S3C) and sensitization to the DSB-inducing antibiotic ciprofloxacin (Figs 5 and S2), except for the *rdgC* deletion.

Pol I is known to utilize its polymerase activity to fill ssDNA gaps during Okazaki fragment synthesis and following DNA repair [49, 51, 60, 61]. The results shown here suggest this activity is especially important in Δ*priB* cells (Figs 2B and 4B). We hypothesize that persistent ssDNA gaps are formed in *polA12(ts)* mutant strains at elevated temperatures, which could lead to DSBs if left unrepaired (Fig 8). This notion is supported by *polA12(ts)* Δ*priB* phenotype suppression on minimal media (Fig 4B) when multi-fork DNA replication is less likely to occur and cause DSBs from collisions with ssDNA gaps [28-30].

The formation of DSBs in *lexA* or *rdgC* deletion strains is less straightforward. Previous work has shown that the absence of PriA or Rep helicase activity can allow the RecFOR mediator proteins to inappropriately load RecA at stalled replication forks [40, 45]. Upon binding, RecA can physically reverse the stalled fork forming a “chicken-foot” structure (Fig 8). DSBs will form from these structures when they are encountered by subsequent replication forks or when they are processed by RuvABC (Fig 8) [28, 63, 66]. Therefore, we hypothesized that the higher levels of DSBs formed in *lexA* or *rdgC* mutants (Figs 6B and S3A,B,D) was caused by excessive RecA activity: either by disrupting its transcriptional repressor (LexA) or by removing a RecA inhibitor (RdgC). Increasing the activity of RecA by disrupting *lexA* or *rdgC* would in turn promote unwarranted RecA activity (Fig 8). Consistent with this notion, we were able to suppress the effects of *lexA* or *rdgC* mutations on Δ*priB* cells by disabling the RecFOR pathway (with a *recR* deletion) or by inhibiting cellular RecA activity by decreasing its expression with a promoter mutation (P_*recA*_(AtoG)) (Fig 7). Notably, these suppression attempts significantly restored the growth rates of Δ*rdgC* Δ*priB* colonies, while permitting (albeit limited) viability of *lexA::kan* Δ*priB* cells. Therefore, it is likely that the SOS DNA-damage response induces the expression of one or more genes (other than *recA*) that are harmful to Δ*priB* cells.

DSBs can form in a variety of different ways in the cell. Disrupting genes identified in the Δ*priB Tn*-seq screen likely increased DSB levels by promoting the formation of DSB-prone substrates (stalled/reversed replication forks and ssDNA gaps), which are encountered by subsequent replication complexes in rich media (Fig 8) [28-30]. While DSBs are problematic, cells can survive if they are readily recognized and repaired. In *E. coli*, DSB repair is usually carried out by RecBCD, which processes DSBs before loading RecA to catalyze strand invasion and create a D-loop site for DNA replication restart (Fig 8) [73]. The DSBs formed in *rep, lexA, polA, dam*, and *rdgC* mutants can still be recognized and repaired by the RecBCD pathway to form D-loops, which subsequently undergo DNA replication restart via the PriA/PriB pathway. We hypothesize that these mutations are synergistic with a *priB* deletion because DSBs are committed to a nonproductive pathway (when *priB* is absent) and stagnant D-loops may ultimately lead to cell death (Fig 8). Furthermore, while most DSB-causing mutants showed some sensitization to ciprofloxacin, *priB* and *recA* deletion strains exhibited extreme sensitization (Fig 5). Taken together, our data support a fundamental link between DSB repair and the PriA/PriB pathway of DNA replication restart.

The results presented here highlight a variety of new questions and exciting opportunities of study. While *uup, nagC*, and *rdgB* are conditionally important in Δ*priB* cells, their disruption does not appear to cause DSBs in the conditions tested (S3A,B,D Fig). Most puzzling is the genetic relationship between *priB* and *nagC*, a transcriptional repressor that coordinates the biosynthesis of *N*-acetylglucosamine, a component of the bacterial cell wall [33, 34]. Deletion of *nagC* led to an aberrant cell morphology (S3A Fig), which may have caused the mutant’s extremely low level of mean fluorescence in our experiments (S3E Fig). It is possible the perturbed cell membrane morphology is linked to DNA damage, similar to observations made with perturbed nuclear envelopes upon loss of lamin proteins in cancer cells [74]. Future studies will be required to further probe this possibility. Taken together, our findings have defined a primary role for the PriA/PriB replication restart pathway following DSB repair in *E. coli* and have established important links that integrate replication restart processes into a larger genome maintenance program in bacteria.

## Materials and Methods

### Strain construction

All strains used in this study are derivatives of *E. coli* MG1655 (S1 Table). To enhance the viability and ease of cloning, all strains (unless otherwise stated in S1 Table) carry the *sulB103* allele, encoding a FtsZ variant that resists SulA-mediated cell division inhibition [26, 27]. All plasmids and oligonucleotides used in this study are listed in S2 Table. To construct derivative *polA12(ts)* and MuGam-GFP strains, the method developed by Datsenko and Wanner [75] was employed with some modifications, as described previously [43]. All strains constructed with P1 transduction utilized kanamycin selection, many of which relied on Keio collection strains as donors [76]. All chromosomal mutations were confirmed with PCR amplification flanking the locus of interest, and if necessary, verified with Sanger sequencing.

### Transposome preparation

Transposon mutagenesis was performed using the EZ-Tn5 <DHFR-1> transposon kit (Epicentre) and EK54/MA56/LP372 Tn5 transposase, a hyperactive variant [77]. The Tn5 transposon was PCR amplified with oAM054 and Phusion polymerase (New England Biolabs). Tn5 transposase was purified as described previously [78, 79]. Transposomes were prepared by incubating 2.5 pmol of Tn5 DNA with 0.5 nmol of Tn5 transposase in 20 µL for 3 hr at room temperature before dialyzing into 1x-TE for 3 hr to remove salt prior to electroporation.

### Generation of electrocompetent cells and *in vivo* transposition

*E. coli* strains were prepared for transposition as previously described [79]. Briefly, cells in mid-log phase were washed three times with ice-cold 10% glycerol. In the final wash, cells were either resuspended in 10% glycerol or glycerol-yeast extract medium, flash frozen with liquid nitrogen, and stored at -80 °C. Dialyzed transposome (5 µL) was mixed with 100 µL of electrocompetent cells, electroporated, and immediately recovered in 1 mL of SOC medium for 1 hr. After recovery, dilutions of the cells were plated on SOB-agar containing 10 µg/mL trimethoprim to select for transposon-insertion mutants. Colony counts for each library were estimated by counting one-third of ∼10% of plates. To pool the mutants and construct libraries of ∼500,000 insertion mutants, 2 mL of LB was added to each plate to scrape the colonies into a thick slurry. Care was taken to sufficiently mix each slurry before archiving each in technical triplicate (in 50% glycerol) at -80 °C.

### Preparation of transposon-insertion DNA for sequencing

For sufficient sampling, 100 mL of LB (with 10 µg/mL trimethoprim) was inoculated to OD_600_ ∼0.02 with each respective transposon-insertion mutant library and grown overnight at 37 °C. Genomic DNA was purified using a Wizard Genomic DNA Purification Kit (Promega) and quantified using the QuantiFluor ONE dsDNA System (Promega). Genomic DNA was sheared to ∼200 bp fragments with sonication. The resulting gDNA fragments were prepared for sequencing using NEBNext Ultra II DNA Library Prep Kit for Illumina (New England Biolabs). Bead-based size selection was used to enrich for 200 bp fragments prior to a 21-cycle splinkerette PCR utilizing a custom Tn5-enriching forward primer (oAM055) and custom indexed reverse primers for multiplexing (oAMrev) [25]. To ensure the quality and length of amplified DNA, a final bead-based size selection was employed. DNA was then sequenced with a NextSeq platform (Illumina) at the University of Michigan Advanced Genomics Core using a custom read primer (oAM058) to read the last 10 nt of the transposon before entering chromosomal DNA (to ensure reads corresponded to Tn5-insertions). To maintain sufficient sequence diversity on the flow-cell, a phiX174 DNA spike (20%) was also included in the run. A custom index read primer (oAM059) and standard Illumina primer (oAM112) were employed for sequencing the read indexes and PhiX DNA, respectively.

### *Tn*-seq data analysis

*Tn*-seq sequencing files were trimmed with fastx_trimmer.pl version 0.0.13.2 <u>(htt</u>p<u>://hannonlab.cshl.edu/fastx_toolkit)</u> using default parameters except the first base to keep (-f flag) was set to 10 to remove transposon sequence. Individual samples were then split with fastx_barcode_splitter.pl, version 0.0.13.2 <u>(htt</u>p<u>://hannonlab.cshl.edu/fastx_toolkit)</u> using a file containing the sample ID and the individual barcode sequence used to split each sample into an individual FASTQ file. The barcode sequence was then removed from each read within each FASTQ file using Cutadapt, version 1.13 [80]. The trimmed FASTQ files were then aligned to the *E. coli* K-12 MG1655 genome (NC_000913.3) using Bowtie2, version 1.2 using default parameters [81]. Conditionally important or essential genes were determined using TSAS, version 0.3.0 using Analysis_type2 for two-sample analysis to compare transposon insertion profiles of each mutant strain to the *wt* [31]. Weighted read ratios were calculated as described previously [31]. All other parameters were kept at the default settings.

### Plasmid (*priB*-pRC7) retention assay

The *priB*-pRC7 plasmid is a lac^+^ mini-F (low copy) derivative of pFZY1 [42] containing the *priB* gene. PCR amplification of *priB* with oAM170 and oAM171 conferred ApaI restriction sites flanking the gene. The resulting PCR product and the empty pRC7 plasmid were digested with ApaI and ligated, yielding *priB*-pRC7. Gene deletions via P1 transduction were carried out after the cells had been transformed with the *priB*-pRC7 plasmid to help ensure the viability of each mutant tested. Once constructed, cultures were grown overnight in LB supplemented with 50 µg/mL ampicillin. The following day, cells were diluted 100x in LB and grown to ∼0.2 OD_600_ shaking at 37 °C. The cultures were then placed at 4 °C, serially diluted, and plated on SOB-agar containing X-gal (80 µg/mL) and IPTG (1 mM) to yield 50-500 colonies per plate. Most colonies were counted and imaged after 16 hr incubations at 37 °C, but plates used in Fig 7 were incubated for 22 hr to better visualize the small white colonies.

### Growth competitions

A growth competition experiment was used to determine if deleting *rdgB* conferred a measurable fitness defect in Δ*priB* cells. Pairwise competitions were constructed where the fitness effect of a Δ*rdgB* mutation was examined in a *priB*^+^ or Δ*priB* strain. To quantify the abundance of the Δ*rdgB* mutant, one strain within each competition was modified to carry a neutral Δ*araBAD* mutation. When *ara*^-^ or *ara*^+^ strains are plated on medium containing tetrazolium and arabinose, they form red or white colonies, respectively. The individual strains of each competition were grown in isolation overnight at 37 °C in LB, then equivalent volumes of each were mixed and diluted 100x in fresh LB. The cultures (now with competing strains) resumed growth at 37 °C, and incubations were temporarily paused every 24 hr to re-dilute (100x) in fresh LB and quantify the Δ*rdgB* mutant abundance by plating on LB-agar with tetrazolium (0.005% w/v) and arabinose (1% w/v). The competitions were performed in biological triplicate and with pairwise alternation of the Δ*araBAD* mutation (to ensure it did not produce a fitness effect).

### Spot plating experiments

Serial dilution spot plating was used to examine mutant sensitivities to ciprofloxacin and the effect of temperature and media on *polA12(ts)* strains. For ciprofloxacin sensitivity experiments, biological triplicate LB cultures were inoculated and grown overnight at 37 °C, whereas strains used in the *polA12(ts)* experiment were grown at 30 °C. The following day, the cultures were diluted to OD_600_ of 1.0 and 10x serial dilutions were prepared with the corresponding plating media (LB or M9). Serial dilutions (10 µL) ranging from 10^−1^ to 10^−6^ were spot plated and incubated at 37 °C, unless stated otherwise. LB-agar plates were incubated for 16 hr, and M9-agar plates were incubated for 40 hr before imaging.

### Fluorescence and brightfield microscopy

An *E. coli* strain carrying MuGam-GFP (SMR14334 [56]) was derivatized to carry the *sulB103* allele (*wt*) before P1 transduction deleted other genes of interest. Saturated cultures were diluted 100x and grown in LB for 30 min at 37 °C to enter early exponential phase. MuGam-GFP expression was then induced at 100 ng/mL doxycycline and growth continued for an additional 2.5 hr at 37 °C. Cells were pelleted and resuspended in 1x PBS buffer (to OD_600_ of 1.0) and placed on ice. About 15 min prior to imaging, cell membrane stain FM 4-64 (5 mM) was added and 2-3 µL of cells were sandwiched between a 24×50 mM, No. 1.5 coverslip (Azer Scientific) and a 1.5% agarose pad. All cells were imaged at room temperature with a motorized inverted Nikon Ti-eclipse N-STORM microscope equipped with a 100x objective and ORCA Flash 4.0 digital CMOS C13440 (Hamatsu). Imaging was performed using NIS-Elements software with the microscope in epifluorescence mode. Cells were first imaged in the brightfield (4.5 V, 100 ms exposure). Visualization of the cell membranes was performed in the DsRed channel to ensure the focusing (4.5 V, 50 ms exposure) and then MuGam-GFP was imaged in the GFP channel (4.5 V, 50 ms exposure). Growth, preparation, and imaging was performed for each strain in biological triplicate.

Analysis of cell features was performed with Fiji software (ImageJ) equipped with plugins as described previously: Single Molecule Biophysics (https://github.com/SingleMolecule/smb-plugins) and MicrobeJ [82]. Briefly, the nd2 raw images for each strain (4 to 8 per replicate with a maximum difference of 2 images within triplicate) were concatenated together by channels. The image processing of each channel was carried out the same way and uniformly throughout the field of view. The scale of all images was corrected to fit the Hamamatsu camera scale. The brightfield and DsRed image stacks were auto-scaled while the GFP images were processed with discoidal averaging of 1-5 and intensity scale set at 0-300. Both brightfield and DsRed channels were cleaned by running a Bandpass filter 10_2 with autoscale 5, a rolling sliding stack of 10, and an enhance contrast of 0.1. Channel stacks were converted to 8 bits before analysis in MicrobeJ. For the analysis, hyperstacks combining only the FM 4-64 and GFP channels were generated in MicrobeJ. From these hyperstacks, cell outlines were detected in the DsRed channel using the default method with a threshold of +25. Within identified cells, GFP foci were detected using the maxima features as foci with a Gaussian fit constraint. The exact setup used to identify bacteria and MuGam-GFP foci in MicrobeJ is available (Final Bacteria setup 1_5 foci 90) as a .xml file. After automatic detection, cells were manually sorted to remove poorly fitting outlines or outlines fitting to cells out of focus. Cell features analysis acquired with MicrobeJ (cell ID, cell length, number of foci per cell, foci intensity and size) were exported as .csv files. Plots and statistical analysis were generated and performed with GraphPad Prism software. At least 650 single cells were analyzed for each condition.

## Acknowledgements

The authors thank Elizabeth Wood, Michael Cox, Steven Sandler, and Susan Rosenberg for generous donations of strains used in this study. We thank Steven Sandler, Jade Wang, Melissa Harrison, and members of the Keck laboratory for critical reading of the manuscript. This study made use of UW-Madison’s Biochemistry Optical Core, and the authors thank Peter Favreau for technical support. This study also made use of UW-Madison’s Biotechnology Center and University of Michigan’s Advanced Genomics Core.

## Supporting Information Captions

**S1 Fig. *Tn*-seq performed in Δ*priB, priC::kan*, and *priA300 E. coli* strains**. (A) Volcano plot (generated with VolcaNoseR) for *Tn*-seq results in Δ*priB* cells. The fold-change (log_2_) in unique insertions within each gene (in *wt* vs Δ*priB* strains) is plotted against the probability of essentiality p-value adjusted for multiple comparisons (-log_10_) [31]. Genes exceeding the fold-change (>4) and significance (>4) thresholds are colored blue and labeled. The *rdgB* gene is also labeled. Circos plots depicting the results of the *Tn*-seq screens in (B) *priC::kan* and (C) *priA300* cells. Each bar in the Circos plots represents the weighted read (log_10_) ratio of a single gene where extension into the blue or orange region corresponds to a detrimental or beneficial, respectively, effect of gene disruption. The *priA300* strain very poorly tolerated transposon insertions within *rep*. Genes with less than three average unique transposon-insertions per replicate in the *wt* condition were omitted.

**S2 Fig. Effects of mutations on DNA damage sensitivity in *E. coli***. The results contained in Fig 4 are expanded to reflect the viabilities of *priC, rdgC, rdgB, uup*, and *nagC* mutants plated on LB-agar with 0-30 ng/mL ciprofloxacin. A *recA* deletion strain was utilized as a positive control of ciprofloxacin hypersensitivity. Dilutions (from left to right) are 10x serial dilutions from normalized overnight culture. Displayed spot plate data are representative of three replicates.

**S3 Fig. DSB formation in mutant *E. coli* strains**. (A) Representative images depicting MuGam-GFP (green) foci and FM 4-64-stained membranes (red) for SMR14334, *wt*, Δ*rep, lexA::kan, polA12(ts)*, Δ*dam*, Δ*priC*, Δ*rdgC*, Δ*uup*, Δ*nagC*, and Δ*rdgB* strains. The brightness of FM 4-64 and GFP images was uniformly exaggerated to highlight differences between strains more clearly. Scale bars are 10 µm. The abundance of MuGam-GFP foci per cell (B), measured cell lengths (C), distribution of the number of foci per cell in cells with MuGam-GFP foci (D), and mean fluorescence per cell (E) are shown for all strains included in A. (B-D) Mean values are depicted with error bars representing standard error of the mean. (E) Median values are depicted as gray or black bars. (B-C) Statistical significance (U-Mann-Whitney) for each strain compared to the *wt* control is displayed: P < 0.05 (*), P < 0.01 (**), P < 0.001 (***), and P < 0.0001 (****).

**S4 Fig. Importance of Pol I polymerase activity in Δ*priB E. coli***. Biological replicates (A,B) of spot plating experiments pertaining to Fig 4B.

**S5 Fig. Mutant sensitivities to ciprofloxacin**. Biological replicates (A,B) of spot plating experiments pertaining to Figs 5 and S2.

**S1 Table. Strains used in this study**.

**S2 Table. Oligonucleotides and plasmids used in this study**. The “*” indicates phosphorothioate bonds. The underlined bases in oAMrev reflect twelve distinct indexes (and primers) employed for multiplexing during sequencing. Italicized bases in oAM192/oAM193 and oAM215/oAM216 directed FRTkanFRT chromosomal insertion location to construct AM354 and AM395 strains, respectively.

**S1 File. Analysis of Δ*priB, priC::kan*, and *priA300 Tn*-seq data**.

**S2 File. Colony counts for *priB*-pRC7 retention assays and growth competitions**.

**S3 File. Fluorescence and brightfield microscopy data/analysis**.

## References

1. Cox MM, Goodman MF, Kreuzer KN, Sherratt DJ, Sandler SJ, Marians KJ. The importance of repairing stalled replication forks. Nature. 2000;404(6773):37–41.

2. Mangiameli SM, Merrikh CN, Wiggins PA, Merrikh H. Transcription leads to pervasive replisome instability in bacteria. eLife. 2017;6.

3. Windgassen TA, Wessel SR, Bhattacharyya B, Keck JL. Mechanisms of bacterial DNA replication restart. Nucleic acids research. 2018;46(2):504–19.

4. Michel B, Sandler SJ. Replication Restart in Bacteria. Journal of bacteriology. 2017;199(13).

5. Sandler SJ. Multiple genetic pathways for restarting DNA replication forks in Escherichia coli K-12. Genetics. 2000;155(2):487–97.

6. Sandler SJ, Leroux M, Windgassen TA, Keck JL. Escherichia coli K-12 has two distinguishable PriA-PriB replication restart pathways. Molecular microbiology. 2021;116(4):1140–50.

7. Lee EH, Kornberg A. Replication deficiencies in priA mutants of Escherichia coli lacking the primosomal replication n’ protein. Proceedings of the National Academy of Sciences of the United States of America. 1991;88(8):3029–32.

8. Nurse P, Zavitz KH, Marians KJ. Inactivation of the Escherichia coli priA DNA replication protein induces the SOS response. Journal of bacteriology. 1991;173(21):6686–93.

9. Masai H, Asai T, Kubota Y, Arai K, Kogoma T. Escherichia coli PriA protein is essential for inducible and constitutive stable DNA replication. The EMBO journal. 1994;13(22):5338–45.

10. McCool JD, Ford CC, Sandler SJ. A dnaT mutant with phenotypes similar to those of a priA2::kan mutant in Escherichia coli K-12. Genetics. 2004;167(2):569–78.

11. Sandler SJ, McCool JD, Do TT, Johansen RU. PriA mutations that affect PriA-PriC function during replication restart. Molecular microbiology. 2001;41(3):697–704.

12. Tougu K, Peng H, Marians KJ. Identification of a domain of Escherichia coli primase required for functional interaction with the DnaB helicase at the replication fork. The Journal of biological chemistry. 1994;269(6):4675–82.

13. Kim S, Dallmann HG, McHenry CS, Marians KJ. Coupling of a replicative polymerase and helicase: a tau-DnaB interaction mediates rapid replication fork movement. Cell. 1996;84(4):643–50.

14. Kim S, Dallmann HG, McHenry CS, Marians KJ. Tau protects beta in the leading-strand polymerase complex at the replication fork. The Journal of biological chemistry. 1996;271(8):4315–8.

15. Costa A, Hood IV, Berger JM. Mechanisms for initiating cellular DNA replication. Annual review of biochemistry. 2013;82:25–54.

16. Sandler SJ, Marians KJ, Zavitz KH, Coutu J, Parent MA, Clark AJ. dnaC mutations suppress defects in DNA replication- and recombination-associated functions in priB and priC double mutants in Escherichia coli K-12. Molecular microbiology. 1999;34(1):91–101.

17. Flores MJ, Ehrlich SD, Michel B. Primosome assembly requirement for replication restart in the Escherichia coli holDG10 replication mutant. Molecular microbiology. 2002;44(3):783–92.

18. Boonsombat R, Yeh SP, Milne A, Sandler SJ. A novel dnaC mutation that suppresses priB rep mutant phenotypes in Escherichia coli K-12. Molecular microbiology. 2006;60(4):973–83.

19. Marinus MG. Recombination is essential for viability of an Escherichia coli dam (DNA adenine methyltransferase) mutant. Journal of bacteriology. 2000;182(2):463–8.

20. Nowosielska A, Marinus MG. Cisplatin induces DNA double-strand break formation in Escherichia coli dam mutants. DNA repair. 2005;4(7):773–81.

21. Heller RC, Marians KJ. The disposition of nascent strands at stalled replication forks dictates the pathway of replisome loading during restart. Molecular cell. 2005;17(5):733–43.

22. Langridge GC, Phan MD, Turner DJ, Perkins TT, Parts L, Haase J, et al. Simultaneous assay of every Salmonella Typhi gene using one million transposon mutants. Genome research. 2009;19(12):2308–16.

23. van Opijnen T, Bodi KL, Camilli A. Tn-seq: high-throughput parallel sequencing for fitness and genetic interaction studies in microorganisms. Nature methods. 2009;6(10):767–72.

24. van Opijnen T, Camilli A. Transposon insertion sequencing: a new tool for systems-level analysis of microorganisms. Nature reviews Microbiology. 2013;11(7):435–42.

25. Barquist L, Mayho M, Cummins C, Cain AK, Boinett CJ, Page AJ, et al. The TraDIS toolkit: sequencing and analysis for dense transposon mutant libraries. Bioinformatics (Oxford, England). 2016;32(7):1109–11.

26. Bi E, Lutkenhaus J. Analysis of ftsZ mutations that confer resistance to the cell division inhibitor SulA (SfiA). Journal of bacteriology. 1990;172(10):5602–9.

27. McCool JD, Long E, Petrosino JF, Sandler HA, Rosenberg SM, Sandler SJ. Measurement of SOS expression in individual Escherichia coli K-12 cells using fluorescence microscopy. Molecular microbiology. 2004;53(5):1343–57.

28. Withers HL, Bernander R. Characterization of dnaC2 and dnaC28 mutants by flow cytometry. Journal of bacteriology. 1998;180(7):1624–31.

29. Fossum S, Crooke E, Skarstad K. Organization of sister origins and replisomes during multifork DNA replication in Escherichia coli. The EMBO journal. 2007;26(21):4514–22.

30. Hill NS, Kadoya R, Chattoraj DK, Levin PA. Cell size and the initiation of DNA replication in bacteria. PLoS genetics. 2012;8(3):e1002549.

31. Burger BT, Imam S, Scarborough MJ, Noguera DR, Donohue TJ. Combining Genome-Scale Experimental and Computational Methods To Identify Essential Genes in Rhodobacter sphaeroides. mSystems. 2017;2(3).

32. Drees JC, Chitteni-Pattu S, McCaslin DR, Inman RB, Cox MM. Inhibition of RecA protein function by the RdgC protein from Escherichia coli. The Journal of biological chemistry. 2006;281(8):4708–17.

33. Pennetier C, Domínguez-Ramírez L, Plumbridge J. Different regions of Mlc and NagC, homologous transcriptional repressors controlling expression of the glucose and N-acetylglucosamine phosphotransferase systems in Escherichia coli, are required for inducer signal recognition. Molecular microbiology. 2008;67(2):364–77.

34. Plumbridge J. DNA binding sites for the Mlc and NagC proteins: regulation of nagE, encoding the N-acetylglucosamine-specific transporter in Escherichia coli. Nucleic acids research. 2001;29(2):506–14.

35. Murat D, Bance P, Callebaut I, Dassa E. ATP hydrolysis is essential for the function of the Uup ATP-binding cassette ATPase in precise excision of transposons. The Journal of biological chemistry. 2006;281(10):6850–9.

36. Romero ZJ, Armstrong TJ, Henrikus SS, Chen SH, Glass DJ, Ferrazzoli AE, et al. Frequent template switching in postreplication gaps: suppression of deleterious consequences by the Escherichia coli Uup and RadD proteins. Nucleic acids research. 2020;48(1):212–30.

37. Bradshaw JS, Kuzminov A. RdgB acts to avoid chromosome fragmentation in Escherichia coli. Molecular microbiology. 2003;48(6):1711–25.

38. Savic DJ, Jankovic M, Kostic T. Cellular role of DNA polymerase I. Journal of basic microbiology. 1990;30(10):769–84.

39. d’Ari R. The SOS system. Biochimie. 1985;67(3-4):343–7.

40. Mahdi AA, Buckman C, Harris L, Lloyd RG. Rep and PriA helicase activities prevent RecA from provoking unnecessary recombination during replication fork repair. Genes & development. 2006;20(15):2135–47.

41. Goodall ECA, Robinson A, Johnston IG, Jabbari S, Turner KA, Cunningham AF, et al. The Essential Genome of Escherichia coli K-12. mBio. 2018;9(1).

42. Bernhardt TG, de Boer PA. Screening for synthetic lethal mutants in Escherichia coli and identification of EnvC (YibP) as a periplasmic septal ring factor with murein hydrolase activity. Molecular microbiology. 2004;52(5):1255–69.

43. Romero ZJ, Chen SH, Armstrong T, Wood EA, van Oijen A, Robinson A, et al. Resolving Toxic DNA repair intermediates in every E. coli replication cycle: critical roles for RecG, Uup and RadD. Nucleic acids research. 2020;48(15):8445–60.

44. Giese KC, Michalowski CB, Little JW. RecA-dependent cleavage of LexA dimers. Journal of molecular biology. 2008;377(1):148–61.

45. Moore T, McGlynn P, Ngo HP, Sharples GJ, Lloyd RG. The RdgC protein of Escherichia coli binds DNA and counters a toxic effect of RecFOR in strains lacking the replication restart protein PriA. The EMBO journal. 2003;22(3):735–45.

46. Joyce CM, Grindley ND. Method for determining whether a gene of Escherichia coli is essential: application to the polA gene. Journal of bacteriology. 1984;158(2):636–43.

47. Joyce CM, Fujii DM, Laks HS, Hughes CM, Grindley ND. Genetic mapping and DNA sequence analysis of mutations in the polA gene of Escherichia coli. Journal of molecular biology. 1985;186(2):283–93.

48. Klenow H, Henningsen I. Selective elimination of the exonuclease activity of the deoxyribonucleic acid polymerase from Escherichia coli B by limited proteolysis. Proceedings of the National Academy of Sciences of the United States of America. 1970;65(1):168–75.

49. Lehman IR, Chien JR. Persistence of deoxyribonucleic acid polymerase I and its 5’--3’ exonuclease activity in PolA mutants of Escherichia coli K12. The Journal of biological chemistry. 1973;248(22):7717–23.

50. Camps M, Loeb LA. Critical role of R-loops in processing replication blocks. Frontiers in bioscience : a journal and virtual library. 2005;10:689–98.

51. Uyemura D, Lehman IR. Biochemical characterization of mutant forms of DNA polymerase I from Escherichia coli. I. The polA12 mutation. The Journal of biological chemistry. 1976;251(13):4078–84.

52. Kogoma T. Stable DNA replication: interplay between DNA replication, homologous recombination, and transcription. Microbiology and molecular biology reviews : MMBR. 1997;61(2):212–38.

53. Willmott CJ, Maxwell A. A single point mutation in the DNA gyrase A protein greatly reduces binding of fluoroquinolones to the gyrase-DNA complex. Antimicrobial agents and chemotherapy. 1993;37(1):126–7.

54. Tamayo M, Santiso R, Gosalvez J, Bou G, Fernández JL. Rapid assessment of the effect of ciprofloxacin on chromosomal DNA from Escherichia coli using an in situ DNA fragmentation assay. BMC microbiology. 2009;9:69.

55. Klitgaard RN, Jana B, Guardabassi L, Nielsen KL, Løbner-Olesen A. DNA Damage Repair and Drug Efflux as Potential Targets for Reversing Low or Intermediate Ciprofloxacin Resistance in E. coli K-12. Frontiers in microbiology. 2018;9:1438.

56. Shee C, Cox BD, Gu F, Luengas EM, Joshi MC, Chiu LY, et al. Engineered proteins detect spontaneous DNA breakage in human and bacterial cells. eLife. 2013;2:e01222.

57. Huisman O, D’Ari R. An inducible DNA replication-cell division coupling mechanism in E. coli. Nature. 1981;290(5809):797–9.

58. Bhattacharyya S, Soniat MM, Walker D, Jang S, Finkelstein IJ, Harshey RM. Phage Mu Gam protein promotes NHEJ in concert with Escherichia coli ligase. Proceedings of the National Academy of Sciences of the United States of America. 2018;115(50):E11614–e22.

59. Mojas N, Lopes M, Jiricny J. Mismatch repair-dependent processing of methylation damage gives rise to persistent single-stranded gaps in newly replicated DNA. Genes & development. 2007;21(24):3342–55.

60. Cao Y, Kogoma T. The mechanism of recA polA lethality: suppression by RecA-independent recombination repair activated by the lexA(Def) mutation in Escherichia coli. Genetics. 1995;139(4):1483–94.

61. Glickman BW. The role of DNA polymerase I in excision-repair. Basic life sciences. 1975;5a:213–8.

62. Michel B, Ehrlich SD, Uzest M. DNA double-strand breaks caused by replication arrest. The EMBO journal. 1997;16(2):430–8.

63. Seigneur M, Bidnenko V, Ehrlich SD, Michel B. RuvAB acts at arrested replication forks. Cell. 1998;95(3):419–30.

64. Robu ME, Inman RB, Cox MM. RecA protein promotes the regression of stalled replication forks in vitro. Proceedings of the National Academy of Sciences of the United States of America. 2001;98(15):8211–8.

65. Courcelle J, Donaldson JR, Chow KH, Courcelle CT. DNA damage-induced replication fork regression and processing in Escherichia coli. Science (New York, NY). 2003;299(5609):1064–7.

66. Seigneur M, Ehrlich SD, Michel B. RuvABC-dependent double-strand breaks in dnaBts mutants require recA. Molecular microbiology. 2000;38(3):565–74.

67. Weisemann JM, Weinstock GM. Direct selection of mutations reducing transcription or translation of the recA gene of Escherichia coli with a recA-lacZ protein fusion. Journal of bacteriology. 1985;163(2):748–55.

68. Weisemann JM, Weinstock GM. The promoter of the recA gene of Escherichia coli. Biochimie. 1991;73(4):457–70.

69. Nguyen B, Shinn MK, Weiland E, Lohman TM. Regulation of E. coli Rep helicase activity by PriC. Journal of molecular biology. 2021;433(15):167072.

70. Syeda AH, Wollman AJM, Hargreaves AL, Howard JAL, Brüning JG, McGlynn P, et al. Single-molecule live cell imaging of Rep reveals the dynamic interplay between an accessory replicative helicase and the replisome. Nucleic acids research. 2019;47(12):6287–98.

71. Heller RC, Marians KJ. Unwinding of the nascent lagging strand by Rep and PriA enables the direct restart of stalled replication forks. The Journal of biological chemistry. 2005;280(40):34143–51.

72. Raghunathan N, Goswami S, Leela JK, Pandiyan A, Gowrishankar J. A new role for Escherichia coli Dam DNA methylase in prevention of aberrant chromosomal replication. Nucleic acids research. 2019;47(11):5698–711.

73. Dillingham MS, Kowalczykowski SC. RecBCD enzyme and the repair of double-stranded DNA breaks. Microbiology and molecular biology reviews : MMBR. 2008;72(4):642-71, Table of Contents.

74. Denais CM, Gilbert RM, Isermann P, McGregor AL, te Lindert M, Weigelin B, et al. Nuclear envelope rupture and repair during cancer cell migration. Science (New York, NY). 2016;352(6283):353–8.

75. Datsenko KA, Wanner BL. One-step inactivation of chromosomal genes in Escherichia coli K-12 using PCR products. Proceedings of the National Academy of Sciences of the United States of America. 2000;97(12):6640–5.

76. Baba T, Ara T, Hasegawa M, Takai Y, Okumura Y, Baba M, et al. Construction of Escherichia coli K-12 in-frame, single-gene knockout mutants: the Keio collection. Molecular systems biology. 2006;2:2006.0008.

77. Goryshin IY, Reznikoff WS. Tn5 in vitro transposition. The Journal of biological chemistry. 1998;273(13):7367–74.

78. Bhasin A, Goryshin IY, Reznikoff WS. Hairpin formation in Tn5 transposition. The Journal of biological chemistry. 1999;274(52):37021–9.

79. Byrne RT, Chen SH, Wood EA, Cabot EL, Cox MM. Escherichia coli genes and pathways involved in surviving extreme exposure to ionizing radiation. Journal of bacteriology. 2014;196(20):3534–45.

80. Martin M. Cutadapt removes adapter sequences from high-throughput sequencing reads. EMBnetjournal. 2011;17(1):3.

81. Langmead B, Salzberg SL. Fast gapped-read alignment with Bowtie 2. Nature methods. 2012;9(4):357–9.

82. Ducret A, Quardokus EM, Brun YV. MicrobeJ, a tool for high throughput bacterial cell detection and quantitative analysis. Nature microbiology. 2016;1(7):16077.

